# Iron and peroxide regulation of the PrrF sRNAs and a conserved Dps-like protein in *Pseudomonas aeruginosa* and *Pseudomonas fluorescens*

**DOI:** 10.1101/2025.11.17.688906

**Authors:** Khady O. Ouattara, Amanda G. Oglesby

## Abstract

*Pseudomonas aeruginosa* is a Gram-negative opportunistic pathogen that requires iron to cause infection. Iron can also be toxic due to its participation in Fenton chemistry, resulting in the production of reactive oxygen species (ROS). Thus, *P. aeruginosa* regulates the uptake, use, and storage of iron to mitigate the effects of ROS. *P. aeruginosa* uses several mechanisms to manage oxidative stress, including superoxide dismutases to detoxify the superoxide radical, catalases to break down H_2_O_2_, and members of the ferritin superfamily to store iron. The iron-responsive PrrF1 and PrrF2 small regulatory RNAs (sRNAs) are predicted to pair with and destabilize mRNA transcripts for several oxidative stress response proteins, including *sodB*, *katA*, and *brnD*, encoding a novel bacterioferritin-like Dps protein. In this study, we developed *brnD* reporter constructs that are responsive to PrrF-mediated iron regulation. We demonstrated that PrrF-mediated regulation of *brnD* occurs in the 5’ untranslated region (UTR) of its mRNA, likely via a conserved region of complementarity with the PrrF sRNAs. We further demonstrated that *brnD* mRNA levels are increased upon hydrogen peroxide treatment. Surprisingly, peroxide treatment also resulted in elevated levels of the PrrF sRNAs, and this induction occurred specifically at the *prrF2* promoter. Lastly, we investigated these regulatory effects in *Pseudomonas fluorescens*, revealing similar iron regulation of PrrF sRNAs and the *brnD* ortholog, as well as peroxide-induced expression of the PrrF sRNAs. Combined, these data highlight a distinct class of iron-responsive Dps-like proteins with potential functional conservation across the pseudomonads, and they reveal a novel aspect of the oxidative stress response involving the PrrF sRNAs.

**IMPORTANCE:** Iron is a crucial micronutrient for *Pseudomonas aeruginosa* survival and virulence, yet it can also be toxic due to production of reactive oxygen species. The PrrF sRNAs are produced in iron limiting conditions and block the expression of several mRNAs involved in *P. aeruginosa’s* oxidative stress response. Here, we make the surprising discovery that oxidative stress induces expression of the PrrF sRNAs. We also provide evidence that PrrF regulation of two mRNA targets is hindered upon oxidative stress. Oxidative stress similarly induced the PrrF sRNAs in *Pseudomonas fluorescens*, indicating conservation of this novel regulatory event. This study therefore highlights a novel and conserved regulatory link between iron homeostasis and oxidative stress protection in the pseudomonads.

## INTRODUCTION

Iron is required for the survival of almost all organisms but can also be toxic due to its reactivity with oxygen. Through Fenton chemistry, the ferrous ion [Fe(II)] catalyzes the formation of reactive oxygen species (ROS), which can react with proteins, lipids, and DNA to cause cellular damage. Aerobic organisms therefore employ numerous proteins to detoxify ROS, including superoxide dismutases, catalases, and alkyl peroxidases (1, 2). Organisms also used members of the ferritin superfamily to protect against the effects of iron-mediated oxidative stress. An important facet of these ferritin-like proteins is their ferroxidase centers that oxidize Fe(II) to the ferric ion [Fe(III)], which is then stored as an insoluble and nonreactive mineral (3, 4). Members of the ferritin superfamily include ferritins (Ftn), bacterioferritins (Bfr), and DNA protection from starved cells (Dps) proteins. Ftns are conserved across all domains of life, while Bfr and Dps proteins are specific to prokaryotes. Ftns and Bfrs both form 24-mers that function primarily as iron storage proteins, and Bfrs are differentiated from Ftns by a methionine residue that coordinates heme (5). In contrast to Bfrs and Ftns, Dps proteins form 12-mers that bind to and protect DNA when the cell is starved for nutrients (6–8). Because of the conservation of ferritin proteins, understanding the mechanisms by which different ferritin superfamily proteins contribute to iron homeostasis may be broadly translated to multiple organisms.

*Pseudomonas aeruginosa* is a ubiquitous environmental bacterium that causes multi-drug-resistant infections in compromised individuals, including those with the hereditary disease cystic fibrosis (CF) (9, 10). *P. aeruginosa* requires iron for growth and infection, yet the mammalian host limits this nutrient during infection through a process referred to as nutritional immunity (11). During infection, *P. aeruginosa* overcomes host-mediated iron limitation through the expression of multiple high affinity iron uptake systems (11). To mitigate the potential for iron-mediated oxidative stress, the expression of these iron uptake systems is regulated by the ferric uptake repressor (Fur), which blocks transcription of these genes when bound to cytosolic iron (12–14). Fur also represses expression of the PrrF1 and PrrF2 small noncoding regulatory RNAs (sRNAs) (15, 16). The PrrF sRNAs mediate an iron sparing response by pairing with and decreasing the stability and translation of mRNAs for non-essential iron-containing proteins (15, 17, 18). The genes for the PrrF1 and PrrF2 sRNAs are located in tandem on the chromosome and share 94% sequence homology. While the PrrF sRNAs appear to be redundant in function (18), the *prrF1* and *prrF2* promoters respond to distinct regulatory signals. Specifically, the PhuS protein binds to the *prrF1* but not the *prrF2* promoter to mediate heme regulation of the PrrF1 sRNA (19, 20), while the two component response regulator AlgR binds to the *prrF2* but not the *prrF1* promoter. The PrrF sRNAs are required for acute murine lung infection, and their expression is conserved in clinical isolates from CF sputa (16, 18). However, our understanding of how the PrrF sRNAs contribute to *P. aeruginosa* virulence remains incomplete.

The PrrF regulon consists of several mRNAs encoding ROS detoxifying and iron storage proteins. Amongst these are SodB, an iron-cofactored superoxide dismutase that detoxifies the superoxide radical into oxygen and H_2_O_2_, and the heme-cofactored catalase, KatA, which converts H_2_O_2_ into water and oxygen (21, 22). PrrF also negatively affects the expression of, and shares complementarity with, an mRNA that is annotated as a probable Bfr (15). The annotation arose from conserved Bfr ferroxidase center residues as well as a potential heme-coordinating methionine residue (5, 23). However, recent structural studies of PA4880 revealed that it complexes as a 12-mer, indicative of a Dps protein, while also possessing a ferroxidase center that is structurally similar to those formed by Bfrs (24). This study further found no evidence of PA4880 interacting with heme (24), and the heme-coordinating methionine residue is not conserved in orthologs from other *Pseudomonas* species (**Fig. S1**). These findings led us to name PA4880 as <u>b</u>acterioferriti<u>n</u>-like <u>D</u>ps protein, or BrnD. Notably, *P. aeruginosa* also encodes a bacterioferritin (BfrB), ferritin (FtnA), and Dps protein (PA0962) (25), yet only BrnD is regulated by PrrF (15, 17).

In this study, we were motivated to understand the impact of the PrrF sRNAs regulation on the expression of oxidative stress genes, with a primary focus on *brnD*, to gain a better understanding of how iron homeostasis and oxidative stress mitigation are linked. We developed *brnD* reporter strains that are responsive to PrrF-mediated iron regulation. Studies of these reporter strains demonstrate that PrrF-mediated regulation of *brnD* occurs in the 5’ untranslated region (UTR) of its mRNA, likely via a conserved region of complementarity with the PrrF sRNAs. PrrF-mediated iron regulation becomes evident during late exponential growth in both shaking and static cultures, the latter of which have been shown to promote microaerobic conditions. We further demonstrate that *brnD* mRNA levels are increased upon hydrogen peroxide treatment. Surprisingly, peroxide treatment also resulted in elevated levels of the PrrF sRNAs, and *prrF1* and *prrF2* transcriptional reporters indicate that this induction is due to increased *prrF2* promoter activity. We further show that PrrF complementarity to the predicted 5’UTR of *brnD* homologues is conserved across pseudomonads. Moreover, we demonstrate that iron represses the PrrF sRNAs and increases *brnD* expression in *Pseudomonas fluorescens* strain Pf0-1, and that peroxide treatment similarly leads to increased PrrF expression in this environmental pseudomonad. Combined, these data demonstrate iron regulation of a conserved Dps-like protein in the pseudomonads. Moreover, this study reveals a novel and conserved response to oxidative stress involving the PrrF sRNAs.

## RESULTS AND DISCUSSION

### Iron activates *brnD* expression in a PrrF-dependent manner

Previous transcriptomic studies show that *brnD* expression is negatively affected by the PrrF sRNAs in iron-depleted conditions (15, 17). To further investigate how PrrF and iron regulate *brnD* expression, we constructed an expression reporter with the *brnD* promoter and 5’ UTR fused to *lacZY* (**Fig. 1A**). The resulting reporter construct was introduced into the CTX phage attachment site on the chromosomes of *P. aeruginosa* PAO1 and an isogenic Δ*prrF* mutant lacking the *prrF1* and *prrF2* genes. Since BrnD is a Dps-like protein and therefore expected to play a role in oxidative stress, which can be exacerbated by high levels of iron, we first aimed to determine the minimum concentration of iron needed to fully de-repress *brnD* reporter activity. For this, the reporter strains were grown in chelexed and dialyzed tryptic soy broth (DTSB) supplemented with FeCl_3_ at various concentrations (0 -100 µM) for 18 h, conditions that have been used for the majority of prior PrrF and iron regulatory studies (16, 26). Analysis of β-galactosidase activity showed that the *brnD* reporter was significantly induced by as little as 20 μM ferric chloride (FeCl_3_) (**Fig. S2A**). Moreover, the Δ*prrF* reporter strain showed significantly increased *brnD* reporter activity in the absence of iron supplementation (**Fig. S2B**), demonstrating that the *brnD* reporter is subject to PrrF-mediated iron regulation.

**Figure 1.**
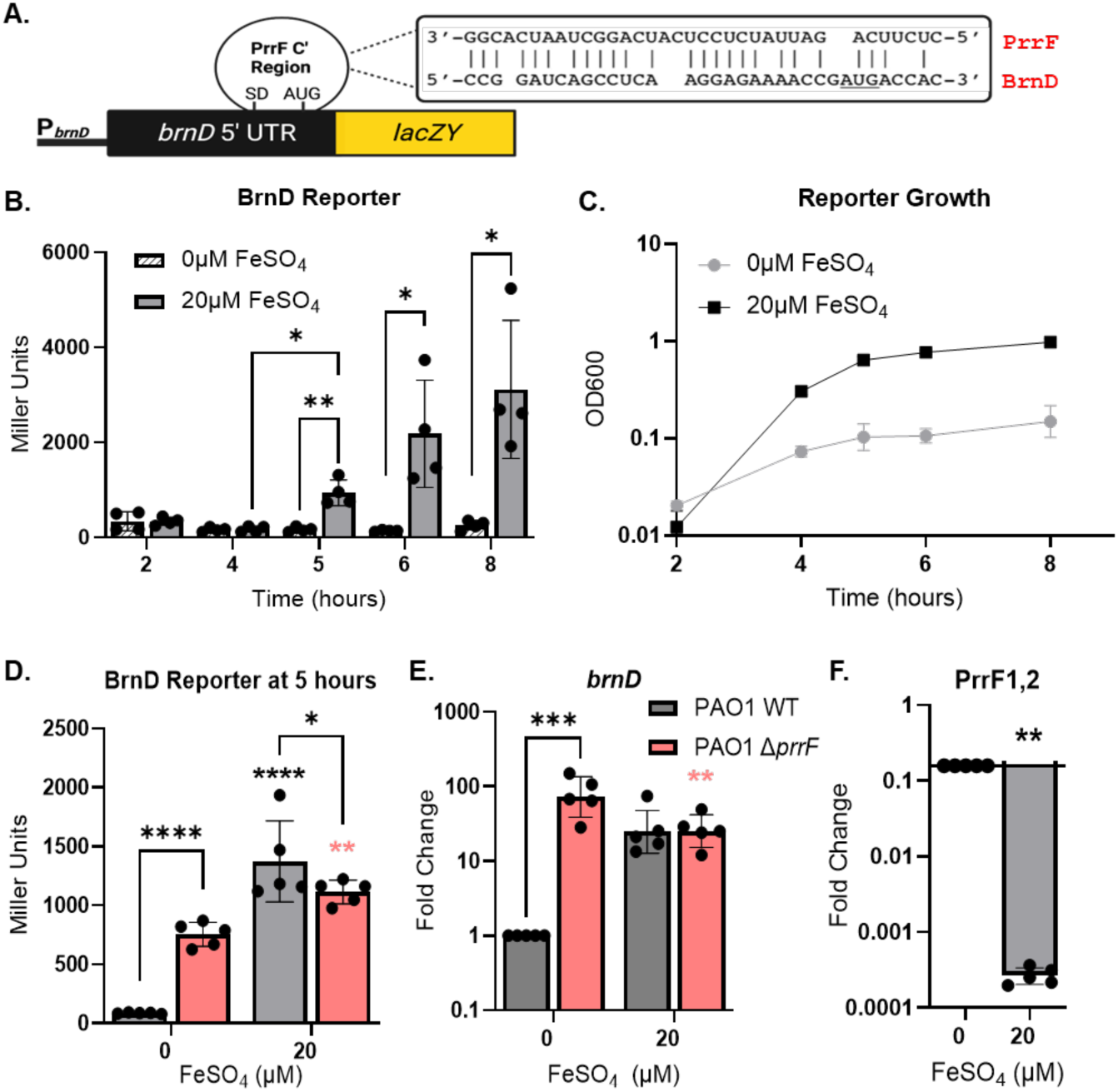
Reporter shows PrrF-mediated iron induction of *brnD*. **A.** Diagram of the *brnD* reporter construct showing the approximate location of complementarity with the PrrF sRNAs overlapping the Shine-Dalgarno (SD) and translational start site (AUG). **B-C.** PAO1 (grey bars) carrying the *brnD* reporter construct was grown with shaking (250 rpm) in CDM with or without supplementation of the indicated concentrations of FeSO_4_ over the span of 8 h and assayed for β-galactosidase activity (**B**) and culture density (**C**). **D-F.** PAO1 and the isogenic Δ*prrF* mutant (pink bars) strains carrying the *brnD* reporter were grown for 5 hours and analyzed for β-galactosidase activity (**D**), and RNA was isolated for qPCR analysis of the *brnD* mRNA (**E**) and the PrrF sRNAs (**F**). Statistics were performed using two-way ANOVA, with Tukey’s multiple comparisons test for significance. Floating asterisks represent strain-specific (black for wild-type and pink for Δ*prrF*) significance in reference to 0 μM FeSO_4_, to which all conditions were normalized: ****P<0.0001, ***P<0.0005, **P<0.005, and *P<0.05.

We next conducted these experiments using a chemically defined medium (CDM) that we have more recently implemented for metal regulation studies (**Fig. S2C-D**) (27, 28). The base CDM is supplemented with 1 mM Ca, 0.3 μM Mn, 6 μM Zn, 0.1 μM Ni, and 0.1 μM Cu, concentrations that have previously been reported in CF sputa (29–31), to avoid mis-metallation in cells upon iron supplementation. We also replaced FeCl_3_ with ferrous sulfate (FeSO_4_) as a soluble iron source to reduce precipitation of the iron salts during culture. Time course analysis of the wild-type *brnD* reporter grown in CDM showed that *brnD* reporter activity is induced by 20 µM FeSO_4_ starting at 5 h of growth (**Fig. 1B**), corresponding with early stationary phase (**Fig. 1C**), and that this induction is lost in the Δ*prrF* mutant reporter strain (**Fig. 1D**). Consistent with analysis of the reporter strains, qPCR showed that *brnD* mRNA levels are similarly increased in the Δ*prrF* mutant compared to wild-type grown in iron-depleted CDM at 5 h of growth (**Fig. 1E**). While *brnD* mRNA levels were 20-fold higher upon iron supplementation of wild-type cultures in CDM, this induction was not statistically significant (**Fig. 1E**). We also analyzed PrrF sRNA expression by qPCR using oligonucleotides that detect both the PrrF1 and PrrF2 sRNAs (18), showing that 20 µM iron significantly downregulates the PrrF RNAs (**Fig. 1F**), correlating with increased *brnD* expression. Since CDM is a defined medium, allowing for better control of metals, we have used this medium for the remainder of the studies described herein.

### PrrF regulation of *brnD* occurs in static culture

Prior expression studies demonstrated that *brnD* is induced upon hypoxia in an Anr-dependent manner (32) and co-regulated with genes for alginate production that support biofilm formation in the hypoxic CF lung environment (33–35). Since our own work has demonstrated that iron and PrrF regulation are altered in hypoxic and anaerobic environments (28, 36), we were prompted to determine whether PrrF-mediated iron regulation of *brnD* would occur in low oxygen environments. To test this idea, we grew the PAO1 wild-type and Δ*prrF* mutant reporter strains in CDM for 12 h in static cultures, which exhibit gene expression profiles indicative of hypoxia (36). We observed significant induction of *brnD* reporter activity with 20 µM iron supplementation after 10 h of static growth (**Fig. 2A**), although we noted that this induction was not consistent from experiment to experiment (**Fig. 2B**), possibly due to overall weaker reporter activity in static versus shaking cultures. However, qPCR analysis showed consistent 2- to 4-fold upregulation of the *brnD* mRNA upon supplementation of cultures with 20 µM iron (**Fig. 2C**). Additionally, *brnD* reporter activity and mRNA levels were significantly de-repressed in the Δ*prrF* mutant compared to the wild-type grown in iron-depleted conditions, indicating that iron induction of *brnD* in the wild-type strain is due to loss of PrrF repression (**Fig. 2B-C**). Also as expected, the PrrF sRNAs were significantly repressed by supplementation of static cultures with 20 µM iron after 10 h of growth (**Fig. 2D**), though not to the same extent as in shaking cultures (**Fig. 1E**). Combined, these data indicate that PrrF-mediated iron regulation of *brnD* occurs in hypoxic conditions.

**Figure 2.**
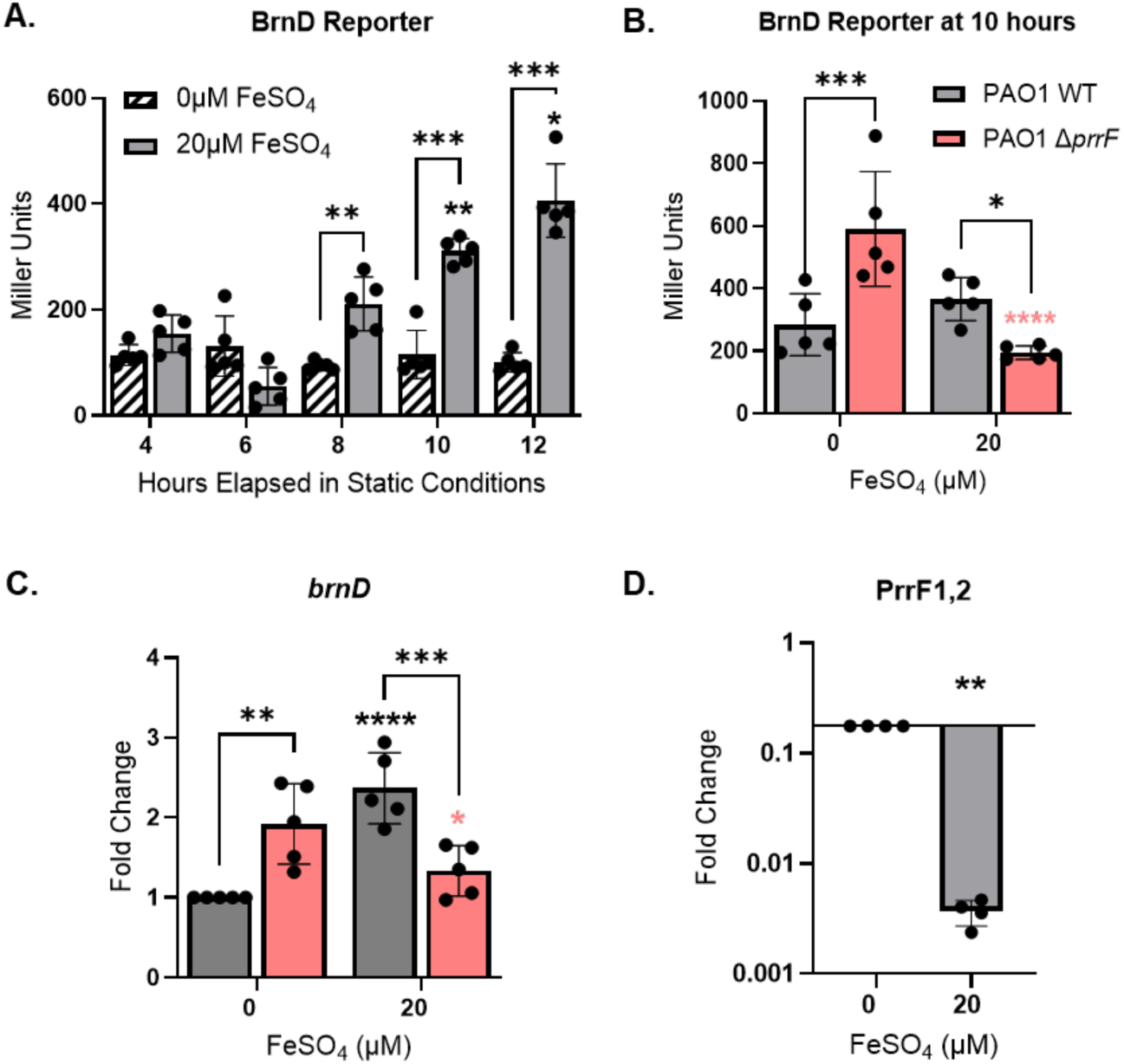
PrrF regulation of *brnD* is conserved in static culture. **A.** PAO1 (grey bars) carrying the *brnD* reporter construct was grown statically in CDM with or without supplementation of 20 µM FeSO_4_ over the span of 12 h and assayed for β-galactosidase activity. **B-D.** PAO1 and the isogenic Δ*prrF* mutant (pink bars) strains carrying the *brnD* reporter were grown statically for 10 h and analyzed for β-galactosidase activity (**B**), and RNA was isolated for qPCR analysis of the *brnD* mRNA (**C**) and the PrrF sRNAs (**D**). Statistics were performed using two-way ANOVA, with Tukey’s multiple comparisons test for significance (**A-B, D**) and one sample t-test for (**C**). Floating asterisks represent condition-specific significance in reference to the 4 h time point (**A**) or strain-specific significance (black asterisks for wild-type and pink asterisks for Δ*prrF*) in reference 0 μM FeSO_4_ (**B-D**): ****P<0.0001, ***P<0.0005, **P<0.005, and *P<0.05.

### PrrF regulation of *brnD* occurs via the 5’ UTR during static growth

To determine if the 5’ UTR of the *brnD* mRNA is responsible for the observed iron and PrrF regulation of *brnD*, we constructed a translational reporter with the *brnD* 5’ UTR fused to the 3’ end of the P*_lac_* promoter and the 5’ end of *gfp* (**Fig. 3A**). The resulting construct was introduced into the CTX phage attachment site of PAO1 wild-type and Δ*prrF* strains. While we did not observe robust iron or PrrF-mediated regulation of fluorescence in shaking cultures (**Fig. S3**), iron-induced fluorescence of the strain was observed after 10 h of static growth (**Fig. 3B**). Moreover, reporter activity was de-repressed in the Δ*prrF* mutant when grown statically in low iron conditions (**Fig. 3B**). We suspect that the lack of iron- and PrrF-regulated fluorescence during shaking growth may be due to the robust activity of the P*_lac_* promoter driving GFP expression prior to PrrF being transcribed by the cells. Nevertheless, the static culture data are consistent with a model wherein the PrrF sRNAs pair with the *brnD* mRNA in the 5’ UTR to affect iron regulation of *brnD* expression.

**Figure 3.**
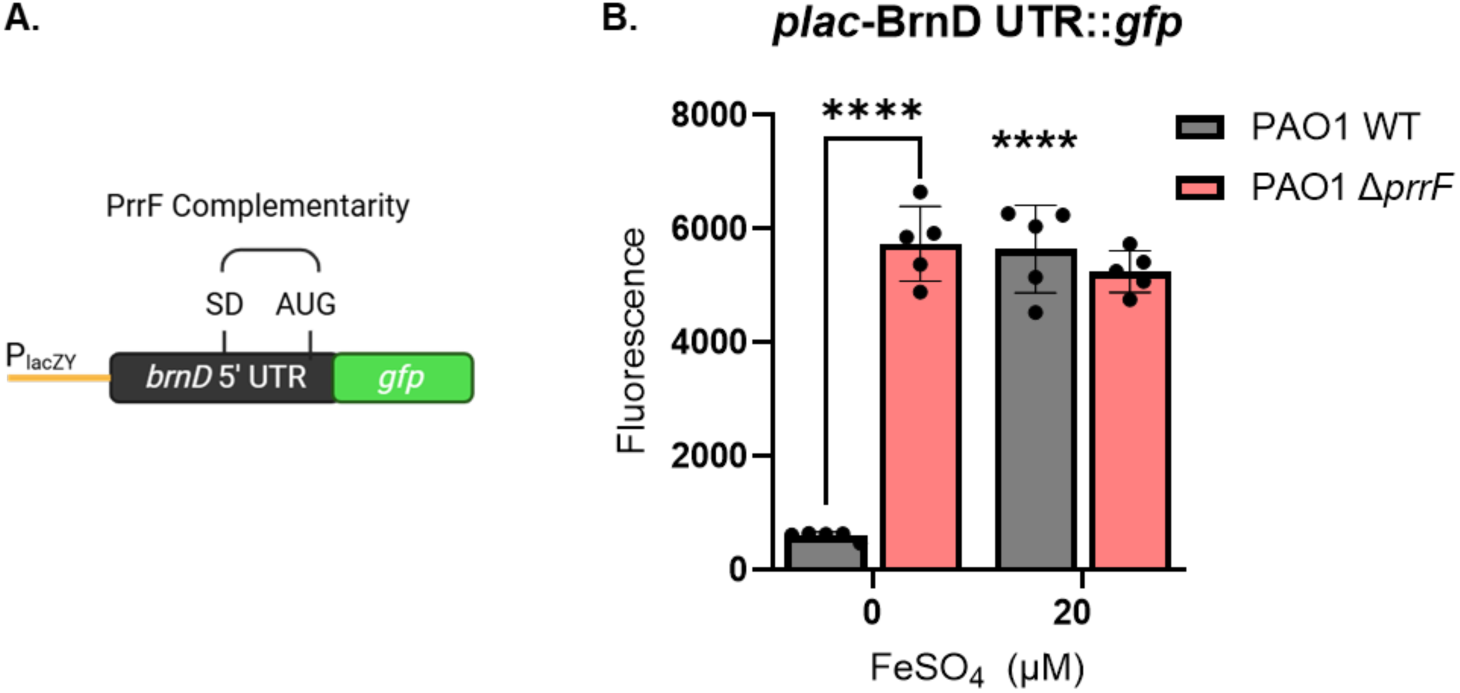
GFP reporter indicates PrrF regulation occurs at brnD’s UTR. **A.** Diagram of the *brnD* translational reporter construct. **B**. Wild-type (gray bars) and Δ*prrF* (pink) strains carrying the *brnD* translational reporter were grown statically for 10 h in CDM with or without supplementation of 20 μM FeSO_4_. GFP was excited at 485nm and fluorescence emitted at 528nm, with a gain of 100. Statistics were done using two-way ANOVA, with uncorrected Fisher’s LSD for significance. Floating asterisks represent significance compared to the WT strain in low iron: ****P<0.0001.

### Peroxide stress increases levels of *brnD* and PrrF RNAs during static growth

Since Dps proteins contribute to oxidative stress protection of many organisms, including *Escherichia coli* and *P. aeruginosa* (8, 25), we next questioned whether *brnD* expression is responsive to oxidative stress. To test this idea, the wild-type and Δ*prrF* strains were grown for 10 h in CDM with 20 µM iron supplementation to promote *brnD* expression, then treated with 5 mM H_2_O_2_. The resulting cultures were analyzed for gene expression by qPCR for 20 minutes post-treatment. Surprisingly, peroxide treatment resulted in significantly increased levels of the PrrF sRNAs, despite cultures being supplemented with iron (**Fig. 4A**). Peroxide treatment also resulted in increased mRNA levels of *brnD* (**Fig. 4B**). As positive controls for peroxide-mediated oxidative stress, we examined expression of *katA* and *sodB*, which were also induced upon peroxide treatment (**Fig. 4C-D**). Induction of *brnD* and *sodB* was prolonged from 10 to 15 minutes post-treatment in the Δ*prrF* mutant compared the wild-type (**Fig. 4B, D**), suggesting that post-transcriptional repression of these two mRNAs by PrrF resumed by 10 minutes post H_2_O_2_ treatment of wild-type cells. In contrast, *katA* mRNA levels remained high in both the wild-type and Δ*prrF* mutant throughout the time course (**Fig. 4C**). Consistent with these data, PrrF negatively affected *sodB* but not *katA* expression during static growth in CDM (**Fig. S4A-B)**. The induction of *sodB* and *brnD* despite increased levels of PrrF suggests that PrrF pairing with these mRNAs is hindered as cells manage their response to oxidative stress. This could be due to another regulatory factor sequestering either PrrF or the target mRNAs, or inactivity of accessory factors that are required for PrrF-mediated gene regulation. For example, host-factor Q (Hfq) is a chaperone protein crucial for the recruitment of the PrrF sRNA and with at least one mRNA target, *antR* (37). Thus, one potential model is that oxidative stress affects Hfq’s activity, resulting in the prolonged increased mRNA levels of *sodB* and *brnD*, even in the presence of PrrF. Alternatively, new transcription of *sodB* and *brnD* upon peroxide treatment may overcome the impacts PrrF-mediated regulation of these mRNAs. Curiously, *katA* expression was unaffected by PrrF in cultures grown statically in CDM, suggesting altered PrrF activity in hypoxic conditions (**Fig. S4B** and **Fig. 4C**). This phenomenon of condition-dependent PrrF activity has previously been observed for the PrrF-responsive *antR* mRNA in aerobic vs anaerobic environments (28) and PrrF regulation of type six secretion proteins in static conditions (36). Understanding how cells mitigate enhanced PrrF expression during oxidative stress is a needed direction for future studies.

**Figure 4.**
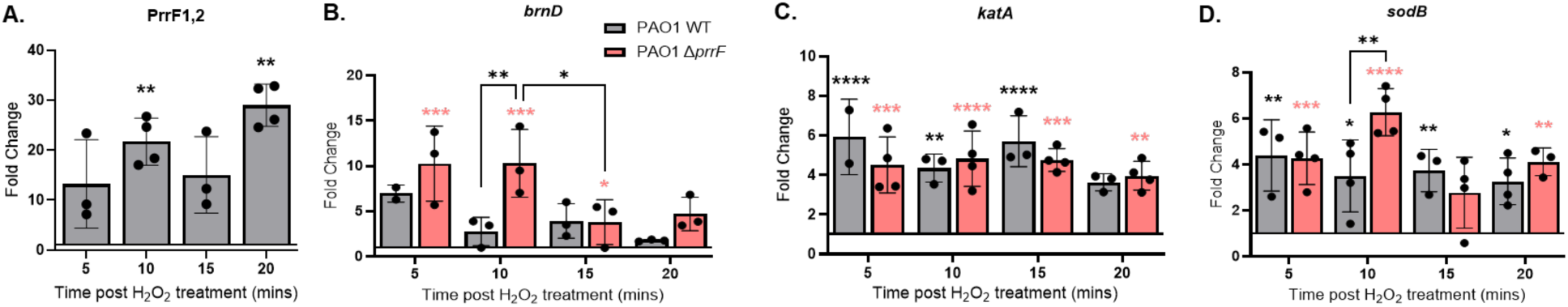
Peroxide stress induces PrrF and its targets in static culture. The indicated strains were grown statically in CDM supplemented with 20 µM FeSO_4_ for 10 h, then treated with 5mM H_2_O_2_ in static conditions, at which point RNA was isolated and used for qPCR analysis of PrrF (**A**), *brnD* (**B**), *katA* (**C**), and *sodB* (**D**). Statistics were done using mixed effects analysis with Dunnett’s multiple comparisons test for (**A**) and two-way ANOVA with Tukey’s multiple comparisons test (**B-D**). Floating asterisks represent strain-specific (black for wild-type and pink for Δ*prrF*) significance in reference to the pre-treatment sample: ****P<0.0001, ***P<0.0005, **P<0.005, and *P<0.05.

### Peroxide treatment results in upregulation of the *prrF2* promoter

The Fur protein binds specifically to Fe(II), which is the predominant ion in the reducing cytosolic environment, to become an active transcriptional repressor (12, 14). A prior study showed that peroxidase and catalase-null *E. coli* mutants exhibited oxidized cytosols, which resulted in deactivation of Fur and de-repression of Fur targets (38). We therefore questioned if increased PrrF sRNA levels upon peroxide treatment was similarly due to Fur inactivation. For this, we determined whether H_2_O_2_ treatment leads to increased transcription initiation using reporter constructs with either the *prrF1* or the *prrF2* promoters fused to *gfp* (**Fig. 5A, D**). The *prrF* reporter strains were grown in CDM supplemented with 20 µM iron for 10 h then treated with 5 mM H_2_O_2_. While we observed a slight increase in both fluorescence and *gfp* mRNA levels upon treatment of the *prrF1* reporter strain, neither of these increases were statistically significant (**Fig. 5B-C**). In contrast, the *prrF2* reporter strain showed significant increases in both fluorescence and *gfp* mRNA levels upon H_2_O_2_ treatment (**Fig. 5E-F**). We interpret the increase in PrrF sRNA levels to be partially due to reduced Fur activity, the apparent specificity of this response for the *prrF2* promoter suggests there are specific attributes of the *prrF2* versus *prrF1* promoter that further contribute to this induction.

**Figure 5.**
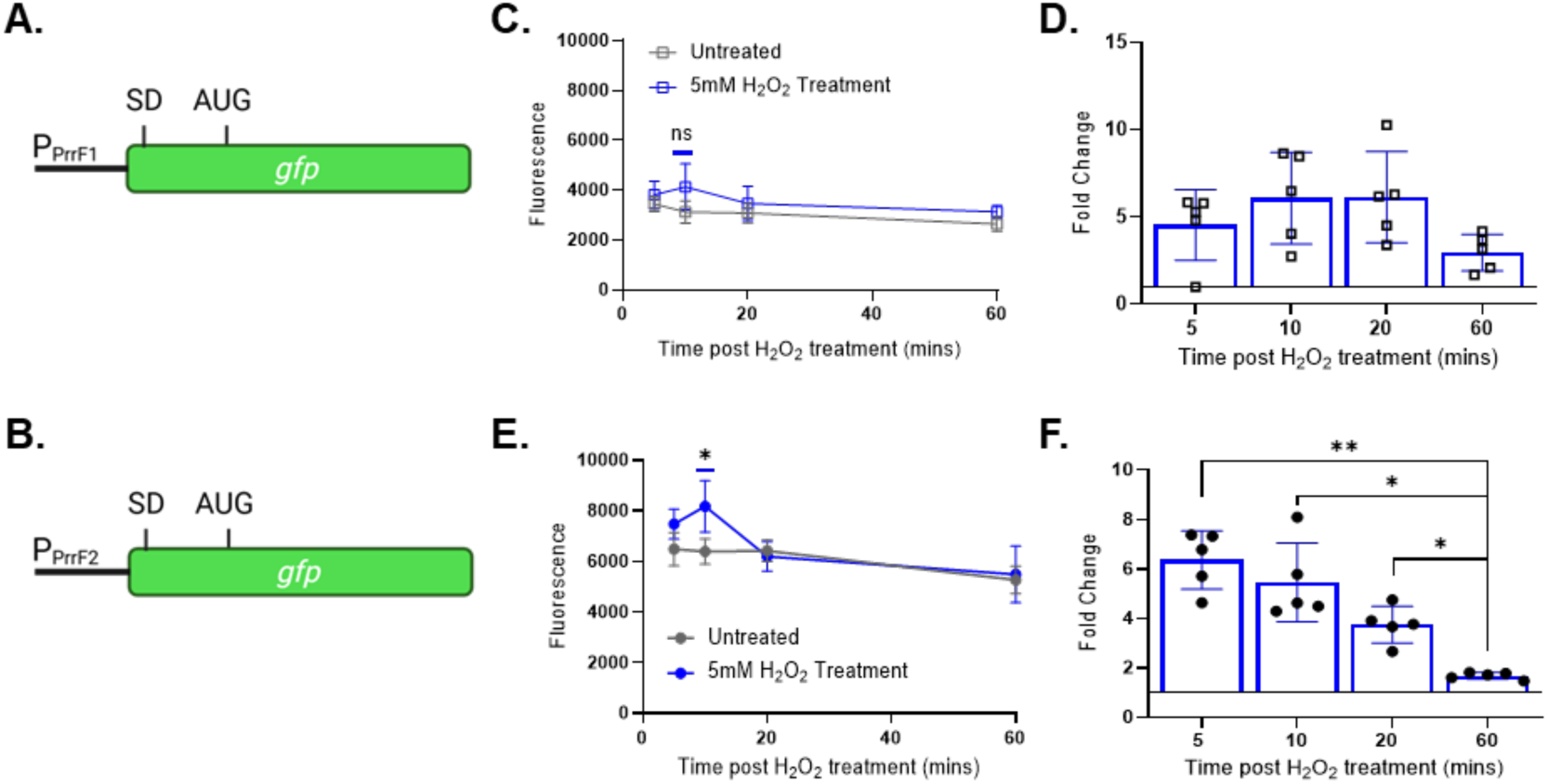
Peroxide differentially induces *prrF2* promoter. **A-B.** Diagrams of the P*_prrF1_* (**A**) and P*_prrF2_* (**B**) transcriptional reporter fusions. SD: Shine Dalgarno. **C-F**. PAO1 carrying the P*_prrF1_* (**C-D**) or P*_prrF2_* (**E-F**) transcriptional reporter constructs were grown statically for 10 h in CDM supplemented with 20μM FeSO_4_ then treated with 5 mM H_2_O_2_. Aliquots were collected and assayed for fluorescence (**C,E**) or harvested for RNA isolation and qPCR analysis of the *gfp* mRNA (**D,F**) at the indicated timepoints post-treatment. GFP was excited at 485nm and fluorescence emitted at 528nm, with a gain of 100. Statistics were performed using two-way ANOVA, with Tukey’s multiple comparisons test for significance: **P<0.005 and *P<0.05.

Previous studies showed that the two component response regulatory AlgR binds to the *prrF2* promoter but not the *prrF1* promoter (39, 40). AlgR is a regulatory protein that binds to the *algD* promoter to activate expression of genes for alginate production, leading to the mucoid phenotype characteristic of chronic *Pseudomonas* infections in the CF lung (41). AlgR has induces several genes involved in oxidative stress protection (42), and the production of alginate has been shown to protect *P. aeruginosa* biofilms against oxidizing agents (43). AlgR also binds to the *pvdS* promoter (39), and we observed a similar significant induction of *pvdS* upon H_2_O_2_ treatment of the *prrF2* transcriptional reporter strain (**Fig. S5**). One intriguing model is that AlgR binding to the *prrF2* promoter upon oxidative stress competes with the more limited pool of active Fur protein, resulting in enhanced *prrF2* promoter activity. The link of AlgR and mucoidy to PrrF2 during oxidative stress reveals yet another layer of regulation on the PrrF sRNAs, and it further suggests the importance of the PrrF2 sRNA in the oxidative stress response of *P. aeruginosa*.

### Iron regulation of *brnD* is conserved in *Pseudomonas fluorescens*

The *brnD* gene is highly conserved in the pseudomonads (**Fig. S1**), and sequence analysis of the *brnD* 5’ UTR and PrrF sRNAs from multiple *Pseudomonas* species shows that PrrF complementarity to the *brnD* mRNA is also broadly conserved (**Fig. 6A**). We therefore wanted to investigate if iron regulation of *brnD* occurs in other pseudomonads. Since the 5’ UTR of *brnD* from *P. fluorescens* strain Pf0-1 is most closely conserved with the 5’ UTR of *brnD* in *P. aeruginosa* PAO1 (**Fig. 6A**), we introduced the *P. aeruginosa brnD* reporter (**Fig. 1A**) into the CTX phage attachment site of *P. fluorescens* Pf0-1. The resulting strain was grown with shaking for 18 h in CDM, and supplementation with either 20 or 100 µM FeSO_4_ resulted in significant induction of β-galactosidase activity (**Fig. 6B**). We similarly observed robust induction of the native *brnD* mRNA in Pf0-1 by qPCR (**Fig. 6C**).

**Figure 6.**
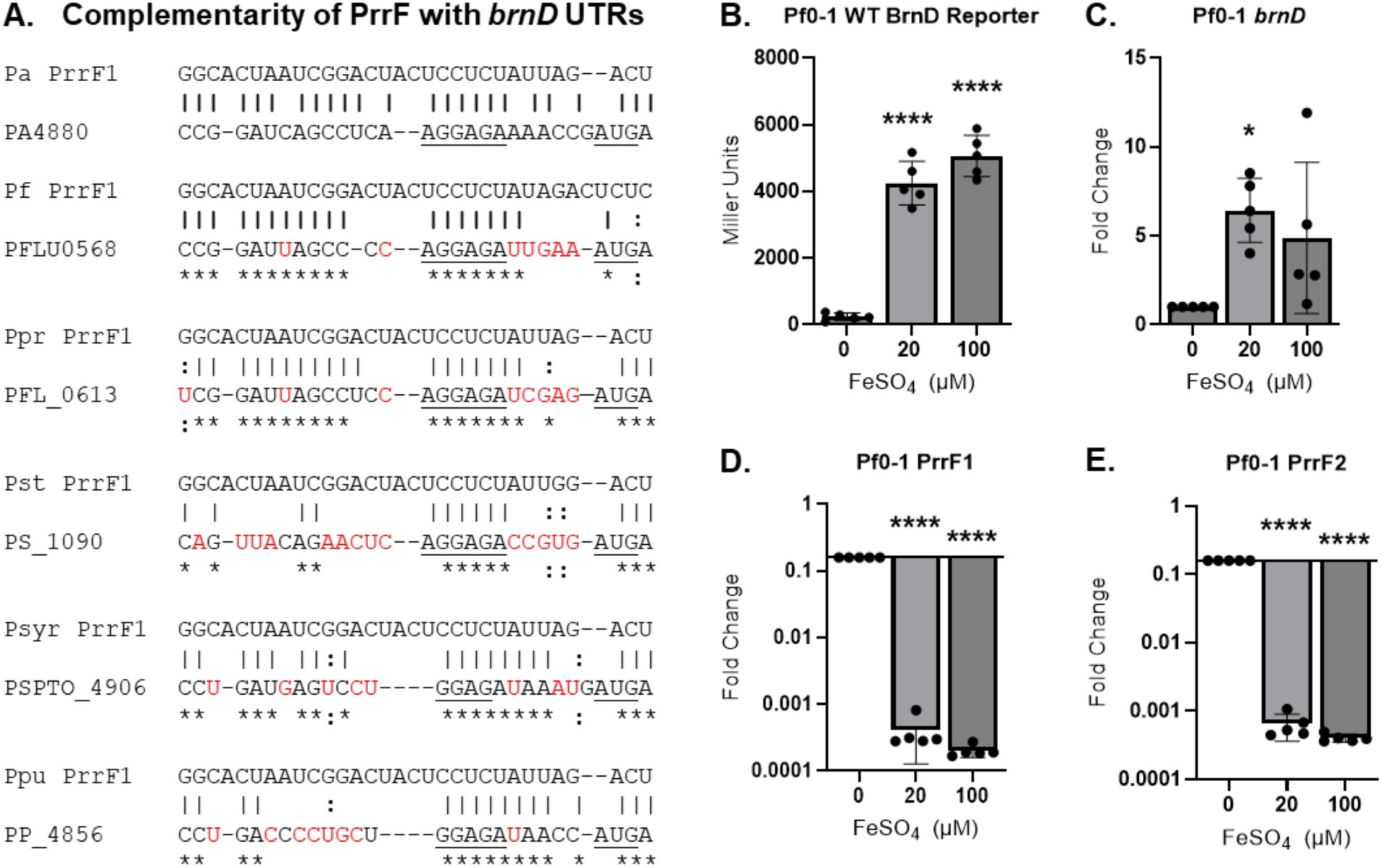
Iron regulation of *brnD* is conserved in Pf0-1. **A**. Complementarity of PrrF homologs in the indicated *Pseudomonas* species with the cognate *brnD* 5’ UTR sequence. **B**. Pf0-1 carrying the *brnD* reporter (**A**) or without (**C-E**) was grown shaking for 18 h in CDM supplemented with or without 20-100 µM FeSO_4_. Cultures were analyzed for β-galactosidase activity (**B**) or harvested for RNA and qPCR analysis of the native *brnD* mRNA (**C**), PrrF1 (**D**), or PrrF2 (**E**) sRNAs. Statistics were performed using One-way ANOVA with Dunnett’s multiple comparisons test for significance: ****P<0.0001 and *P<0.05.

To determine if the increase in *brnD* expression upon iron supplementation correlated with repression of the Pf0-1 PrrF sRNAs, we next examined Pf0-1 PrrF expression by qPCR. Unlike in *P. aeruginosa,* the PrrF1 and PrrF2 sRNAs in *P. fluorescens* are located at distal loci and share only 80% homology (**Fig. S6A**), allowing us to generate primers specific to each PrrF sRNA (**Fig. S6C, Table S2**). Consistent with what we observe in *P. aeruginosa*, iron supplementation significantly repressed both PrrF transcripts (**Fig. 6D-E**). Although iron regulation of *brnD* has previously been observed in this environmental species (44–46), to our knowledge this is the first report showing iron regulation of the PrrF sRNAs in *P. fluorescens*.

### Peroxide stress increases levels of the PrrF sRNAs and the *brnD* reporter in *P. fluorescens*

We next determined whether H_2_O_2_ treatment exerted a similar effect on PrrF and *brnD* expression in Pf0-1 as was observed in *P. aeruginosa*. Time course studies of the Pf0-1 *brnD* reporter strain grown statically in CDM with or without 20 µM iron supplementation for 12 h showed that β-galactosidase activity was induced in iron-replete cultures between 8 and 10 h (**Fig. 7A**), corresponding to late logarithmic phase (**Fig. 7B**). We then grew the Pf0-1 *brnD* reporter strain for 10 h in CDM with 20 µM iron supplementation, treated the cultures with 5 mM H_2_O_2_, and assayed the cultures for β-galactosidase activity at multiple time points post-treatment. Activity of the *brnD* reporter was significantly induced 20 minutes post-treatment as compared to the pre-treatment (**Fig. 7C**), while *brnD* mRNA levels remained unchanged after the addition of peroxide to the cultures (**Fig. 7D**). The disparity between the mRNA and reporter activity results could be attributed to the relatively lower levels of the *brnD* mRNA that we observed in Pf0-1 as compared to PAO1. Alternatively, this finding could be due to differences in how *brnD* is regulated by oxidative stress in *P. fluorescens*. It is also possible that the heterologous *P. aeruginosa brnD* 5’ UTR sequence used to make the *brnD* construct possesses attributes that contribute to peroxide-induced reporter activity in Pf0-1. More studies are needed to understand how *brnD* expression is regulated by oxidative stress in this environmental pseudomonad.

**Figure 7.**
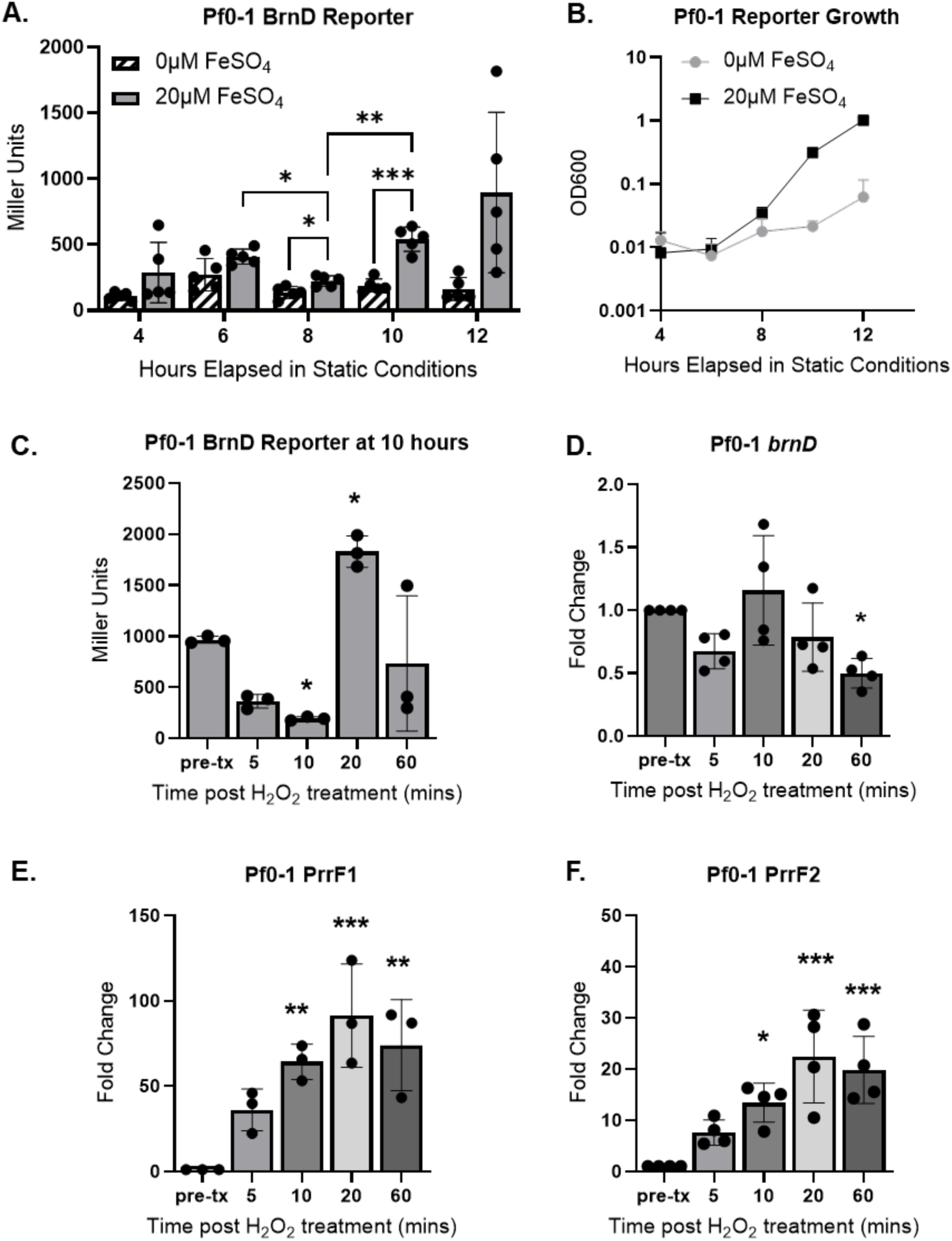
PrrF sRNAs are induced by peroxide treatment in Pf0-1. **A-B.** Pf0-1 carrying the *brnD* reporter was grown statically in CDM supplemented with or without 20 μM FeSO_4_ over a period of 12 h and assayed for β-galactosidase activity (**A**) and culture density (OD_600_) (**B**). **C**. Pf0-1 carrying the *brnD* reporter was grown statically in CDM supplemented with 20 μM FeSO_4_ then treated with 5 mM H_2_O_2_ and assayed for β-galactosidase activity. **D-E**. Cultures of the Pf0-1 strain grown statically and harvested for qPCR analysis of the native *brnD* mRNA (**D**), PrrF1 (**E**), or PrrF2 (**F**) sRNAs. Statistics were performed using two-way ANOVA with Šídák’s multiple comparisons test for significance (**A**) or one-way ANOVA with Dunnett’s multiple comparisons test for significance (**C-F**): ***P<0.0005, **P<0.005, and *P<0.05.

In contrast to the *brnD* mRNA, the PrrF1 and PrrF2 transcript levels were both significantly induced for up to 60 minutes post-peroxide treatment (**Fig. 7E-F**), suggesting this regulatory effect upon peroxide treatment is conserved amongst the pseudomonads. Although *P. fluorescens* is non-pathogenic and does not encounter host-derived oxidative stress, this organism is still susceptible to the deleterious effects of oxidizing agents. Aerobic respiration is one of the main causes of ROS production (14) and auto-oxidation of cellular enzymes can also lead to elevated levels of intracellular peroxide (47, 48). *P. fluorescens* possesses most of the compendium of the alginate regulatory and biosynthetic genes, and alginate aids in *P. fluorescens* adaptation to osmotic stress from drier soil environments (49). Alginate also protects *Pseudomonas putida* biofilms against ROS and osmotic stress (50, 51). Similar to *P. aeruginosa prrF2* (**Fig. S7B**), we have identified a conserved region of that binding site upstream of the *P. fluorescens prrF1* and *prrF2* sequences (**Fig. S7C-D**), extending the AlgR-Fur competition model to this environmental pseudomonad. Future studies into how AlgR affects peroxide-induced expression of the PrrF sRNAs may therefore reveal novel paradigms for oxidative stress responses across the pseudomonads.

### Summary and conclusions

In this study, we investigated the role of the PrrF sRNAs in the regulation of genes involved in oxidative stress protection, with a focus on the recently characterized Dps-like protein BrnD (24). We showed that the PrrF sRNAs negatively affect *brnD* expression and that this regulation is dependent upon the 5’ UTR of the *brnD* mRNA. The expression of *brnD* was also induced by peroxide treatment, despite an increase in PrrF2 transcription and overall increase in PrrF sRNA levels. Notably, this work unveiled yet another differential expression profile in response to peroxide treatment of the two “redundant” PrrF sRNAs, and we present a model wherein AlgR may compete with the Fur protein to affect increase PrrF2 transcription. Combined, these data suggest BrnD plays a role in the oxidative stress response of *P. aeruginosa*, and they present an intriguing new role for the PrrF sRNAs in this response.

Recent structural and functional characterization of BrnD (*aka* DpsL) revealed that this protein has the capacity to store iron in the cavity of the 12-mer Dps fold (24). This work further showed that BrnD, like Dps (PA0962), binds to plasmid DNA *in vitro* (24, 25), suggestive of BrnD playing a role in oxidative stress protection of DNA. Unlike Dps, which contributes to *P. aeruginosa’s* ability to survive H_2_O_2_-mediated oxidative stress (25), an isogenic Δ*brnD* mutant exhibits no phenotype upon H_2_O_2_ treatment in either shaking or static cultures (unpublished data). Given the potential links between BrnD and alginate, we posit that this Dps-like protein plays a more prominent role in biofilms. BrnD also exhibits robust endonuclease activity (24), which could play an important role in restriction protection against the foreign DNA in complex biofilm communities, such as those formed in the CF lung and environment. We are currently pursuing biofilm studies with the Δ*brnD* mutant to understand its role in this important growth condition.

To our knowledge, there is no published work investigating the role of the PrrF sRNAs in other *Pseudomonas* species. In this vein, our current study provides compelling evidence that PrrF-dependent iron regulation of *brnD* is conserved in *P. fluorescens* and throughout the pseudomonads. Moreover, the upregulation of the PrrF sRNAs in response to peroxide treatment is also observed in *P. fluorescens*. Although *P. fluorescens* does not encounter host-derived oxidative stress, this organism is still susceptible to the deleterious effects of oxidizing agents. Aerobic respiration is one of the main causes of ROS production (14), and auto-oxidation of cellular enzymes can also lead to elevated levels of intracellular peroxide (47, 48). However, *P. fluorescens* has evolved to metabolically adapt to respond to oxidative stress insult (52). Like *P. aeruginosa*, *P. fluorescens* possesses genes for BfrB, FtnA, Dps, and BrnD. Bioinformatic analysis using CopraRNA (15) further suggests that *brnD* is the only one of the four ferritin and Dps mRNAs that can pair with PrrF in *P. fluorescens* (unpublished analysis), suggesting this protein imparts an important function for iron homeostasis. More work is needed to understand the how individual pseudomonads have adapted their iron and PrrF regulatory pathways to suit the distinct iron homeostasis needs of different environmental niches.

## MATERIALS AND METHODS

### Bacterial strains, media, and conditions

Strains used in this study are listed in **Table S1**. *P. aeruginosa* was grown in a previously described CDM (53–55). CDM was supplemented with 1 mM CaCl_2_, 0.1 µM CuCl_2_, 0.1 µM NiCl_2_, 6 µM ZnCl_2_, 0.3 µM MnCl_2_ and with or with 20 or 100 µM FeSO_4_, as specified, to afford metal-replete CDM. *P. aeruginosa* lab strains of PAO1 and their deletion mutants were routinely grown overnight by streaking from freezer stocks in tryptic soy agar or brain heart infusion (BHI) agar (Sigma, St. Louis, MO) plates and incubated at 37°C for 12 h–16 h. Three to five colonies were taken from each streaked plate and inoculated in 1.5 mL of tryptic soy broth or LB Broth (Miller) (Sigma, St. Louis, MO), and incubated for 12 h – 16 h with shaking at 250 rpm at 37°C. Shaking cultures were grown aerobically in 5 mL of CDM in 50 mL acid-washed glass flasks with foam stoppers. Cultures for growth assays were inoculated to an OD_600_ of 0.05 and grown for labeled timepoints, with shaking at 250 rpm at 37°C. Static cultures were grown micro-aerobically in 2 mL of high iron, metal-replete CDM in 15 mL plastic culture tubes. Cultures for static growth assays were inoculated to an OD_600_ of 0.05 and grown for ten hours, growing unperturbed in a standing incubator at 37°C. All experiments were performed with at least three biological replicates to ensure reproducibility.

### Generation of reporter strains

To generate the *brnD* translational reporter, the 5’ UTR of *brnD* was amplified by PCR using primers mentioned in **Table S2** (*brnD* UTR For and Rev) and PAO1 genomic DNA as a template. The PCR product was digested by restriction enzymes BamHI and HindIII directly cloned into the multiple cloning site of the P*_lac_*:mini-CTX:GFP^-SD^ vector (**Table S1**). This reporter plasmid lacks the *lacZ* Shine-Dalgarno sequence and was digested by the same restriction enzymes used for the 5’ UTR fragment to generate *P_lac_*:*brnD*-UTR*:gfp^-SD^* plasmid for translational reporter assays. The reporter construct was then introduced into PAO1 and Δ*prrF* chromosomes by integrating them at the CTX *attB* site as previously described (56).

The PAO1 P*_prrF2_*:*gfp* reporter construct was generated in the same manner as the PAO1 P*_prrF1_*:*gfp* reporter (57). Briefly, the promoter region of *prrF2* (P*_prrF2_*) was amplified by PCR using primers in **Table S2** and the PAO1 genomic DNA as a template. The P*_prrF2_* insert was digested with restriction enzymes BamHI and HindIII and ligated into yeast vector pMQ37 [*aka* pLD2477; contains the coding region for GFP (58)] digested with the same restriction enzymes used for the *prrF2* promoter fragment. The ligated pMQ37-P*_prrF2_* plasmid was transformed into *E. coli* strain SM10, then purified, digested with BamHI and EcoRI, and ligated into the mini-CTX-P*_prrF2_*-*gfp* plasmid digested with the same enzymes. The resulting plasmid was transformed in *E. coli* strain SM10, then purified and transformed into *P. aeruginosa* PAO1 via electroporation. The pFLP plasmid (56) was transformed into PAO1 mini-CTX-P*_prrF2_*-*gfp* via electroporation to excise the integrated mini-CTX as previously described (57).

### Beta-galactosidase reporter assays

The *brnD* reporter construct in PAO1 WT and *ΔprrF* backgrounds were assayed for beta-galactosidase activity as previously described with some modifications (59). Briefly, strains were inoculated to an OD_600_ of 0.05 into CDM supplemented with 1 mM CaCl_2_, 0.1 µM CuCl_2_, 0.1 µM NiCl_2_, 6 µM ZnCl_2_ and 0.3 µM MnCl_2_. In addition, 20 or 100 µM FeSO_4_ was supplemented for high iron cultures and grown shaking or static as described above at 37°C. Cells were then aliquoted at listed timepoints and harvested by centrifugation, resuspended in potassium phosphate buffer (30 mM K_2_HPO_4_, 20 mM KH_2_PO_4_), then diluted 1:10 in Z buffer (60 mM Na_2_HPO_4_, 40 mM NaH_2_PO_4_, 10 mM KCl, 1 mM MgSO_4_, and 50 mM β-mercaptoethanol); chloroform and 1% SDS was also added into the reaction mixture. o-nitrophenyl-β-D-galactopyranoside (ONPG) was then used as a substrate to begin the reaction, which was stopped by addition of 1M Na_2_CO_3_, once sufficient yellow color was observed. The OD_420_ was measured for each sample, and the beta-galactosidase activity was calculated using the Miller units formula [(1000 x OD_420_) / (time x volume x OD_600_)].

### GFP reporter assays

Three to five replicates of the *gfp* reporter strains were grown statically for 10 h as described above in either 0, 20, or 100 μM FeSO_4_ supplemented CDM. After 10 h, samples were either treated with or without 5 mM H_2_O_2_, and 200 μL of each sample was aliquoted into black-bottom 96 well plates (Thermo-Fisher Scientific). GFP fluorescence was obtained by exciting samples at 485 nm and emitting at 528 nm, with the laser set to a gain of 100. The OD_600_ readings of the samples were also obtained, to which the fluorescence readings were normalized.

### Real-time PCR

Five cultures of Pf0-1 or PAO1 and the isogenic *ΔprrF* mutant were grown as previously described in CDM, either in shaking or static conditions as indicated. RT-PCR was performed as previously described (18). Briefly, harvested cultures were stored in RNAlater at −80°C. RNA was extracted using the Qiagen RNeasy kit, cDNA was synthesized, and RT-PCR was performed using TaqMan reagents (Roche) and a StepOnePlus system (Thermo-Fisher) for PAO1 strains and SYBR Green reagents and a QuantStudio system (Thermo-Fisher). Standard curves were produced for each primer/primer-probe set listed in **Table S2** in the supplemental material by analyzing cDNA generated from serial dilutions of RNA and used to determine relative amounts of RNAs as described previously (18). Relative RNA levels were then normalized to the levels of 16S ribosomal RNA in respective strains.

### Statistical analysis

Statistical testing is done using Prism 9. Two-way ANOVA with Tukey’s test for multiple comparisons (unless otherwise stated) were used to analyze the statistically significant changes in reporter assays and real-time PCR experiments using at least three biological replicates with a significance threshold of *p* value < 0.05.

## ACKNOWLEDGEMENTS

We thank Ms. Ashley Lykins for generation of the P*_brnD_:*brnD5’UTR:*lacZ* reporter strains and Dr. Tra-My Hoang and Mr. Jake Weiner for generation of the P*_prrF2_*:*gfp* reporter strains. We also thank Ms. Savannah Ford for her assistance generating the Pf0-1 BrnD reporter. Lastly, we thank Prof. George O’Toole for generously sharing the Pf0-1 strain. This work was funded by NIH grants R01AI123320 (to AGO) and R01AI161294 (to AGO).

